# Hybrid female sterility due to cohesin protection errors in oocytes

**DOI:** 10.1101/2025.02.16.638358

**Authors:** Warif El Yakoubi, Bo Pan, Takashi Akera

## Abstract

Hybrid incompatibility can lead to lethality and sterility of F1 hybrids, contributing to speciation. Here we found that female hybrids between *Mus musculus domesticus* and *Mus spicilegus* mice are sterile due to the failure of homologous chromosome separation in oocyte meiosis I, producing aneuploid eggs. This non-separation phenotype was driven by the mis- localization of the cohesin protector, SGO2, along the chromosome arms instead of its typical centromeric enrichment, resulting in cohesin over-protection. The upstream kinase, BUB1, showed a significantly higher activity in hybrid oocytes, explaining SGO2 mis-targeting along the chromosome arm. Higher BUB1 activity was not observed in mitosis, consistent with viable hybrid mice. Cohesion defects were also evident in hybrid mice from another genus, *Peromyscus*, wherein cohesin protection is weakened. Defective cohesion in oocytes is a leading cause of reduced fertility especially with advanced maternal age. Our work provides evidence that a major cause of human infertility may play a positive role in promoting mammalian speciation.

---

Postzygotic reproductive isolation refers to the inability of one population to produce viable and fertile offspring with another population or species. One fundamental goal in evolutionary biology is to identify the mechanisms underlying hybrid incompatibilities, which can act as reproductive isolating barriers (*1–7*). Previous research has identified hybrid incompatibility in chromosome condensation in oocytes from hybrid mice between *Mus musculus domesticus* (hereafter *domesticus*) and *Mus spretus*, leading to female subfertility (*8*–*10*). Chromosome condensation is regulated by the condensin complex, which is one of the SMC (Structural Maintenance of Chromosomes) complexes that regulates chromosome structure and dynamics (*11*, *12*). SMC complexes are ancient enzymes that have evolved significantly in metazoans (*13*), and therefore, we hypothesized that their mis-regulations may represent a common species barrier. To test this idea, we investigated hybrid females between *domesticus* and *Mus spicilegus* (hereafter *spicilegus*), another closely-related species that we can produce viable F1 hybrids in both directions (although we primarily used *spicilegus* as the sire except for fig. S1C) (*14–17*) (Fig. 1A). We identified that *domesticus* x *spicilegus* hybrid females are sterile (Fig. 1B) whereas hybrid males were aggressive, and we were not able to examine their fertility.

**Fig. 1.**
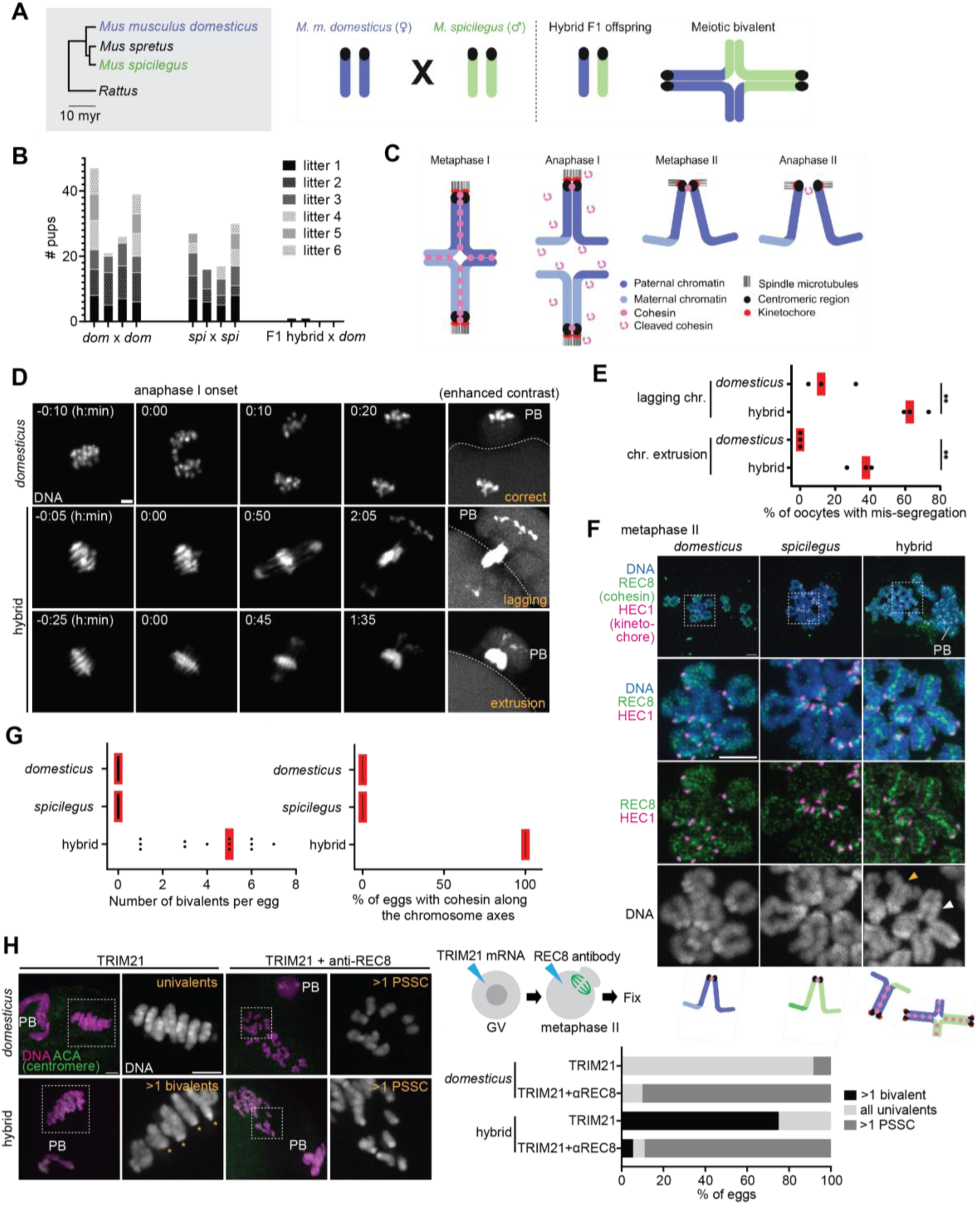
Cohesin mis-regulations lead to non-separation of homologs in hybrid oocytes. (**A**) Phylogenetic tree of mouse species (myr, million years) and schematic of the hybrid mouse system to study hybrid incompatibility in female meiosis. (**B**) 6-month fertility test from indicated genotypes. Age-matched *domesticus* (dom), *spicilegus* (spi) and the hybrid female (F1 hybrid) were used. Each column represents the number of pups per litter for each breeding cage (n = 4 breeding cages per genotype). Note that two F1 female hybrids gave birth to one pup each, which died within few days after the birth. (**C**) Schematic of meiotic chromosome segregation. Cohesin is initially loaded along the chromosome axes and holds sister chromatids together. Cohesin cleavage along the chromosome arm at anaphase I allows the segregation of homologous chromosomes. During metaphase II, sister chromatids maintain their cohesion by the residual cohesin at the pericentromere. Cleavage of this remaining cohesin at anaphase II leads to sister chromatid segregation. (**D**) DNA was visualized in *domesticus* and hybrid oocytes by incubating with SPY-DNA or expressing the Separase biosensor, H2B-mScarlet-Rad21- mNeonGreen (see Fig. 2A) to live-image anaphase I; the mScarlet images are shown in the figure; PB, polar body; dashed lines, oocyte cortex. (**E**) Anaphase chromosome lagging rates in **D** were quantified (n = 68 and 65 oocytes for *domesticus* and hybrid, respectively); red lines, mean; unpaired two-tailed t test was used for statistical analysis; \*\**P* <0.01. (**F**) Chromosome spreads were performed at metaphase II using *domesticus*, *spicilegus* and hybrid oocytes and stained for HEC1 and REC8 (right); orange arrowhead, univalents; white arrowhead, bivalents. (**G**) The number of bivalents per egg and the percentage of eggs with cohesin along the chromosome axes were quantified using the images in **F** (left bottom, n = 13, 13, and 13 eggs for *domesticus*, *spicilegus*, and hybrid); each dot in the graph represents a single egg; red line, median. (**H**) *domesticus* and hybrid oocytes expressing mCherry-Trim21 with or without the anti-REC8 antibody were fixed at metaphase II and stained for ACA (centromere). The percentage of meiosis II eggs with >1 bivalent, >1 precocious separated sister chromatids (PSSC), and normal univalents were quantified (n = 12, 10, 16, and 18 eggs for *domesticus* + TRIM21, *domesticus* + TRIM21 + anti-REC8, hybrid + TRIM21, and hybrid + TRIM21+anti- REC8); scale bars, 5 *µ*m. Schematics in **A** and **C** were created using BioRender.

To understand the mechanism underlying the hybrid female sterility, we imaged chromosome dynamics live from metaphase I to anaphase I in hybrid oocytes (Fig. 1C and 1D). During meiotic prophase, homologous chromosomes pair and recombine to form meiotic bivalents, which segregate at anaphase I to become univalents in meiosis II (*18–21*) (Fig. 1C). The timing of anaphase I onset can be inferred by cell membrane protrusions, which eventually lead to the polar body formation (Fig. 1D, see Fig. 2A for the membrane protrusion at anaphase I onset). In pure *domesticus* oocytes, anaphase I onset coincided with the separation of bivalents (Fig. 1D). In contrast, bivalents failed to segregate properly at anaphase I onset in hybrid oocytes (Fig. 1D and 1E). Mis-segregated chromosomes were either trapped between the egg and the polar body (i.e., lagging chromosomes) or excluded to the polar body (i.e., chromosome extrusion). Mis-segregation in hybrid oocytes was followed by cytokinetic failures, producing polar bodies with variable sizes (fig. S1A). These severe meiotic errors explain why the hybrid female is sterile.

**Fig. 2.**
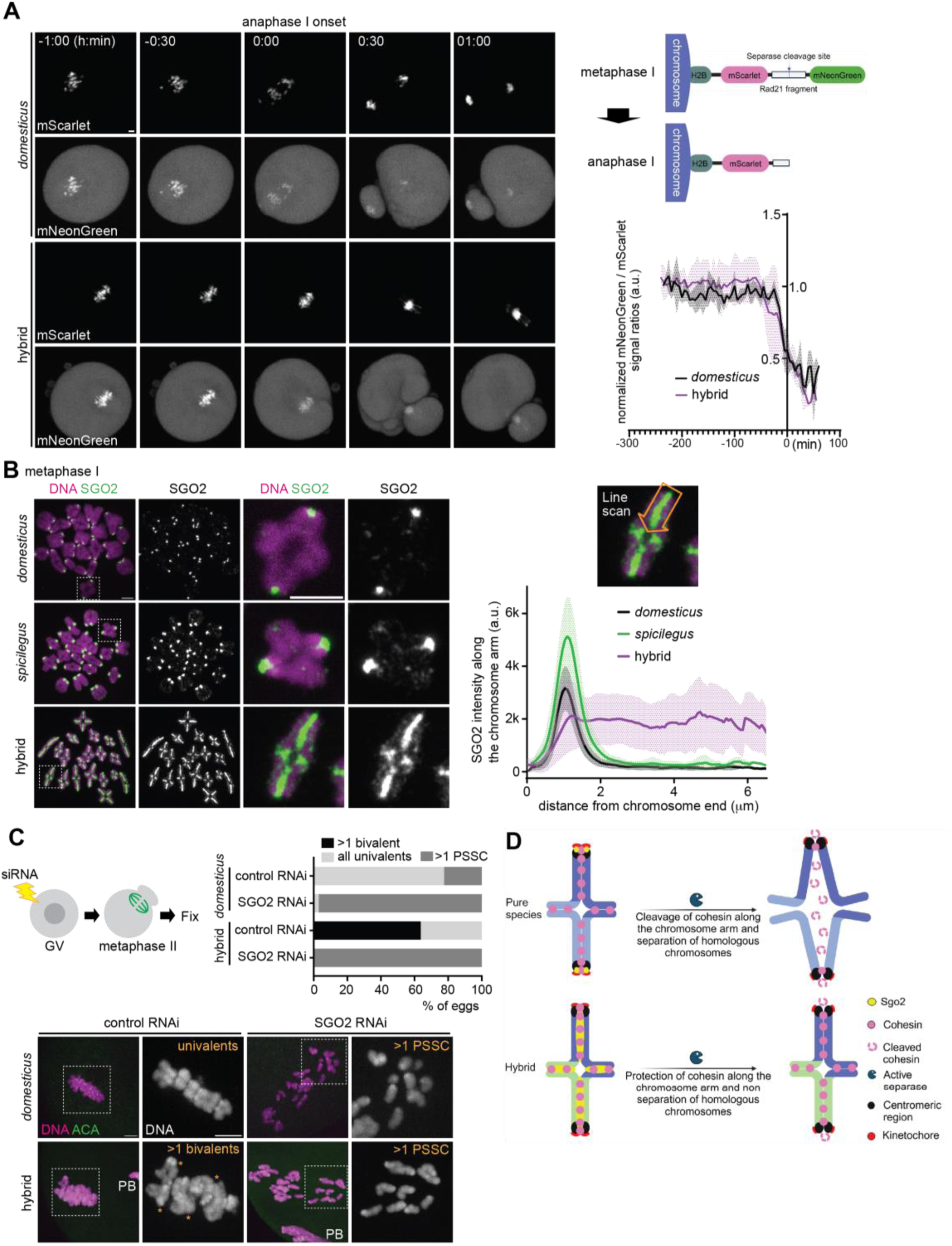
Cohesin over-protection by the ectopic targeting of SGO2. (**A**) Schematic of the Separase biosensor construct (right top). *domesticus* and hybrid oocytes expressing the Separase biosensor were imaged from metaphase I to anaphase I (left). The ratio of mNeonGreen signals divided by mScarlet signals on the chromosomes were quantified over the time course (right bottom, n = 3 and 3 oocytes for *domesticus* and hybrid, respectively); line graph shows the mean values of the mNeonGreen / mScarlet ratio with the shades representing standard deviation. (**B**) Chromosome spreads were performed at metaphase I using *domesticus*, *spicilegus*, and hybrid oocytes and stained for SGO2. Line scans of SGO2 signals were performed along the chromosome arm starting from the chromosome end with a centromere. Line graph shows the mean values of SGO2 intensities along the chromosome arm with the shades representing standard deviation (n = 26, 36, and 60 chromosomes for *domesticus*, *spicilegus*, and hybrid). (**C**) *domesticus* and hybrid oocytes electroporated with control or SGO2 siRNA were fixed at metaphase II and stained with ACA. The percentage of eggs with >1 bivalent, >1 precocious separated sister chromatids (PSSC), and normal univalents were quantified (n = 58, 68, 11, and 17 eggs for *domesticus* + control RNAi, *domesticus* + SGO2 RNAi, hybrid + control RNAi, and hybrid + SGO2 RNAi); scale bars, 5 *µ*m. (**D**) Model of hybrid incompatibility in cohesin protection leading to hybrid female sterility. SGO2 localizes to the centromere and protects cohesin at the pericentromere in pure species oocytes. In contrast, SGO2 ectopically localizes along the arm in hybrid oocytes, protecting cohesin along the arm and causing non-separation of homologous chromosomes; schematics in **D** were created using BioRender.

To understand the cause of this mis-segregation, we analyzed their chromosome morphology in metaphase II eggs. In pure species, chromosomes were all univalents, indicating successful separation of homologous chromosomes at anaphase I (Fig. 1F and 1G). In contrast, hybrid meiosis II eggs carried multiple bivalents (Fig. 1F, hybrid, white arrowhead, Fig. 1G and fig. S1B). Meiosis II eggs with bivalents are aneuploid because the resulting embryos would carry three copies of the chromosome, causing embryonic lethality. Based on these observations, we concluded that defective separation of bivalents at anaphase I in hybrid oocytes leads to chromosome mis-segregation and egg aneuploidy, explaining hybrid female sterility.

We speculated that the defective homolog separation is due to a mis-regulation of cohesin, an evolutionary conserved SMC complex that coheres sister chromatids in mitosis and meiosis (*22–25*). Cohesin undergoes a characteristic two-step removal from the chromosome in meiosis: cohesin is initially loaded along the chromosome arms during meiotic prophase and cleaved by Separase at anaphase I except for the pericentromeric pool, which is critical for bi- orientation of the chromosome on the meiosis II spindle (*26–29*) (Fig. 1C). We analyzed the localization pattern of a meiosis-specific cohesin subunit, REC8, in metaphase II (*22*) (Fig. 1F). REC8 cohesin was localized between sister kinetochores, connecting sister chromatids in pure species as reported previously (*30–32*). Interestingly, REC8 signals were present between sister chromatids along the entire arm of unseparated bivalents in hybrid eggs (Fig. 1F and 1G). Univalents in hybrid eggs also showed similar REC8 distribution along the arm, which is consistent with the observation that sister chromatids of univalents were zippered up unlike typical univalents connected exclusively at the pericentromere (Fig. 1F, hybrid, orange arrowhead). We observed similar REC8 localization and defective homolog separation in hybrid females produced by the reciprocal cross (i.e., *spicilegus* female x *domesticus* male) (fig. S1C), suggesting that the observed phenotypes are not due to maternal or paternal effects. To confirm that defective homolog separation is indeed due to the remaining REC8 on the chromosome, we depleted REC8 by the Trim-Away method (*33*, *34*) (Fig. 1H). REC8 degradation resulted in the separation of chromatids in both *domesticus* and hybrid eggs, indicating that REC8 cohesin holds homologs together in hybrid eggs (rather than DNA catenation etc.).

Aging reduces cohesin levels on the chromosome in oocytes (*35–37*). Therefore, we wondered if reduced cohesin levels in aged oocytes will make the separation of homologs easier in this hybrid. Consistent with this idea, we found that meiosis II eggs from aged hybrid females carried fewer bivalents than those from younger hybrid females (Fig. S2). This observation supports the idea that the non-separation phenotype is caused by the cohesin remaining on the chromosome.

Cohesin on the chromosome arm is phosphorylated during meiosis I, permitting the Separase-mediated cleavage and homologous chromosome segregation at anaphase I (*38–40*). Cohesin at the centromere region is dephosphorylated by the SGO-PP2A complex, protecting cohesin from Separase and maintaining sister-chromatid cohesion until metaphase II (*41–43*). Oocytes with inactive Separase fail to separate homologous chromosomes at anaphase I, proceeding to metaphase II with bivalents (*29*, *44*). Similarly, high SGO-PP2A activity leads to over-protection of cohesin along the arm, also leading to defective separation of homologs (*29*). Based on these studies, we propose two possible models to explain the defective homolog separation in *domesticus* x *spicilegus* hybrid oocytes. First, the Separase activity is lower in the hybrid, causing non-separation of homologs. Second, the SGO-PP2A activity is higher in the hybrid, resulting in cohesin over-protection and defective homolog separation.

To test the first possibility, we used a Separase activity sensor (Fig. 2A), which is an ideal tool to measure the Separase activity during mitosis and meiosis (*45–47*). This sensor construct is composed of a Separase-cleavage site of a cohesin subunit, Rad21, targeted to chromosomes through the N-terminal fusion of histone H2B with two fluorescent proteins, mScarlet and mNeonGreen, flanking the Separase-cleavage site. High Separase activity at anaphase I would remove mNeonGreen from the fusion protein, while mScarlet signals would remain on the chromosome. Live imaging during the metaphase I – anaphase I transition revealed that both *domesticus* and hybrid oocytes showed similar rates of loss of mNeonGreen signals on the chromosomes (Fig. 2A). This result implies that the Separase activity is not significantly affected in hybrid oocytes. Consistent with this observation, sister kinetochores were split in meiosis II hybrid eggs (Fig. S3), which is indicative of active Separase cleaving centromeric cohesin (*29*). Therefore, Separase is active in hybrid oocytes, but is unable to cleave the cohesin on the arm to allow homolog separation.

Next, we tested the second possibility regarding cohesin overprotection by the SGO- PP2A complex. SGO2, the major SGO protein in mouse oocytes (*48*, *49*), localizes to the (peri)centromere region to protect cohesin. Since SGO2 is typically restricted to the centromere, cohesin along the arm is not protected and becomes cleaved by Separase at anaphase I. To test if SGO2 is mis-regulated in hybrid oocytes, causing over-protection, we analyzed the localization pattern of SGO2. We found that SGO2 was aberrantly localized along the chromosome axes in contrast to its centromeric localization in both pure species (Fig. 2B). To test whether the defective separation of homologous chromosomes depends on SGO2, we depleted SGO2 by RNAi (Fig. 2C). In *domesticus* oocytes, depletion of SGO2 during meiosis I caused precocious separation of sister chromatids (PSSC) due to the deprotection of cohesin (*48*, *49*). SGO2 depletion in hybrid oocytes also resulted in PSSC, indicating that defective homolog separation depends on SGO2. Collectively, these results suggest that SGO2 mis-localization drives cohesin protection along the arm, preventing cohesin cleavage at anaphase I and leading to the formation of aneuploid eggs (Fig. 2D).

A conserved kinetochore kinase, BUB1, is essential for the SGO2 localization in mouse and human oocytes by phosphorylating histone H2A (i.e., H2ApT121 in mice) (*32*, *50*, *51*). Since BUB1 is the upstream kinase for SGO2, we wondered if BUB1 mis-regulation causes the ectopic localization of SGO2 in hybrid oocytes. We found that while BUB1 is enriched at the kinetochore in both pure species, BUB1 levels were increased at the kinetochore and also along the chromosome arm in hybrid oocytes (Fig. 3A). As a result, the H2ApT121 mark covered the entire chromosome in hybrid oocytes instead of its typical enrichment near the centromere in pure species (Fig. 3B). We also analyzed the localization of BUB1 and H2ApT121 in mitosis, using granulosa cells (i.e., mitotically growing cells in the ovary). In both hybrid and pure species, BUB1 and H2ApT121 remained restricted to the centromere in mitosis (Fig. 3C), suggesting that the hybrid incompatibility in cohesin protection is specific to (female) meiosis, consistent with viable hybrid mice.

**Fig. 3.**
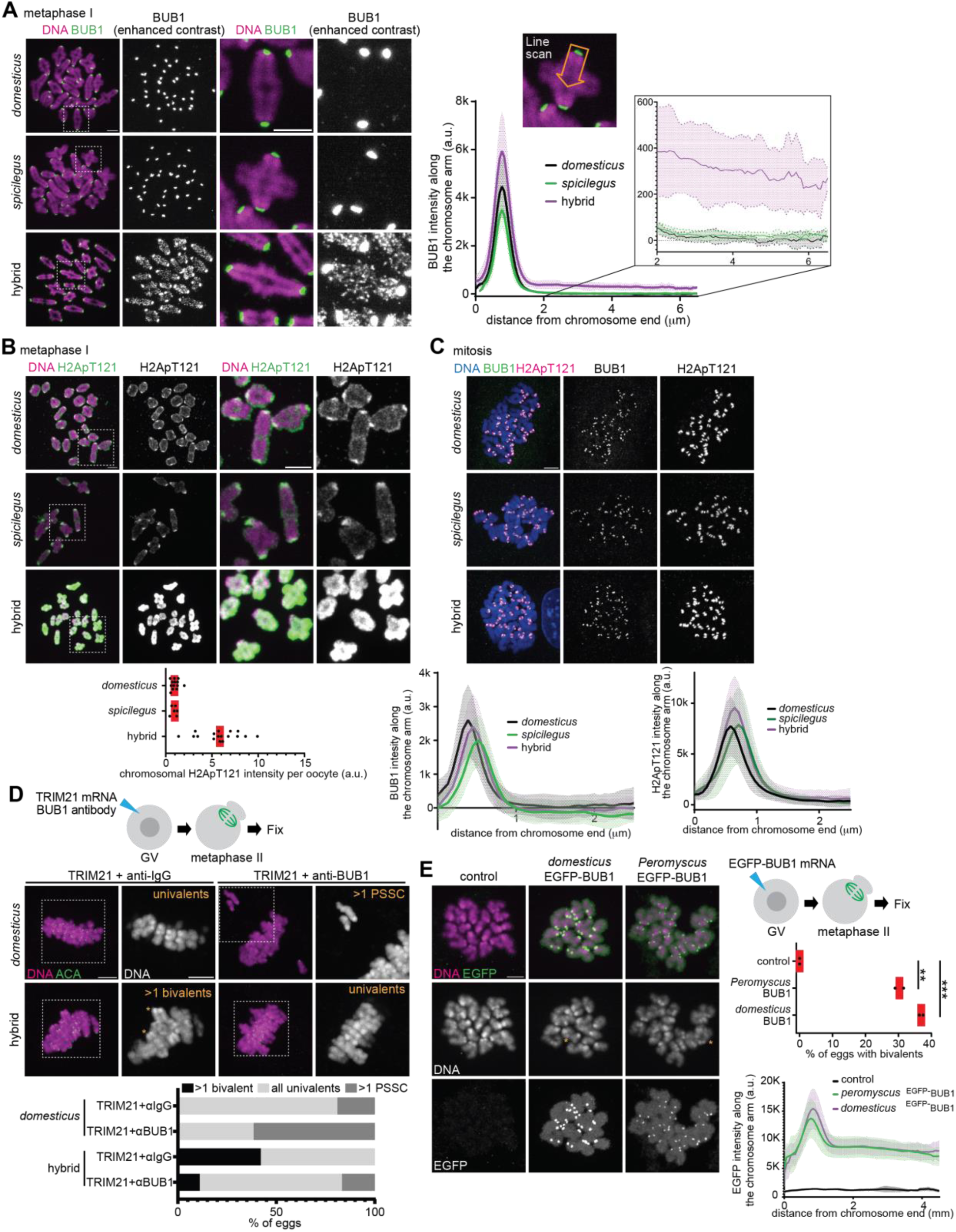
The BUB1-H2ApT121 pathway is up-regulated in hybrid oocytes. (**A**) Chromosome spreads were performed at metaphase I using *domesticus*, *spicilegus*, and hybrid oocytes and stained for BUB1. Line scans were performed to quantify BUB1 signal intensities along the chromosome arm starting from the centromere. Line graph shows the mean values of BUB1 intensity along the chromosome arm with the shades representing standard deviation (n = 26, 36, and 60 chromosomes for *domesticus*, *spicilegus*, and hybrid). The enlarged area of the graph highlights the difference of BUB1 intensity along the chromosome arm. (**B**) Chromosome spreads were performed at metaphase I using *domesticus*, *spicilegus*, and hybrid oocytes and stained for H2ApT121. Graph is the quantification of chromosomal H2ApT121 signal intensities per oocyte. (n = 13, 6, and 17 oocytes for *domesticus*, *spicilegus*, and hybrid); each dot in the graph represents a single oocyte; red line, median. (**C**) Ovarian granulosa cells from *domesticus*, *spicilegus*, and hybrid were fixed and stained for BUB1 and H2ApT121. Line graph is the quantification of BUB1 and H2ApT121 signal intensities along the chromosome arm starting from the centromere. Lines indicate the mean values of BUB1 and H2ApT121 intensities, and the shades represents standard deviation (n = 64, 93, and 102 chromosomes for *domesticus*, *spicilegus*, and hybrid). (**D**) *domesticus* and hybrid oocytes expressing mCherry-TRIM21 with the control IgG or the anti-BUB1 antibodies were fixed at metaphase II and stained for ACA. The percentage of eggs with >1 bivalent, >1 precocious separated sister chromatids (PSSC), and normal univalents were quantified (n = 21, 26, 38, and 54 eggs for *domesticus* + TRIM21 + IgG, *domesticus* + TRIM21 + anti-BUB1, hybrid + TRIM21 + IgG, and hybrid + TRIM21 + anti- BUB1). (**E**) *domesticus* oocytes expressing EGFP-BUB1 derived from *domesticus* or *Peromyscus maniculatus* were fixed at metaphase II and stained for EGFP. The percentage of meiosis II eggs with >1 bivalent was quantified (n = 31, 26, 39 oocytes for control, *domesticus* EGFP-BUB1, *Peromyscus* EGFP-BUB1, respectively); each dot in the graph represents a single oocyte; red line, mean; unpaired two-tailed t test was used for statistical analysis; \*\**P* <0.01, \*\**\*P* <0.001; scale bars, 5 *µ*m.

To directly test if the higher BUB1 level causes the cohesin protection error in hybrid oocytes, we depleted BUB1 by Trim-Away (Fig. 3D). In control *domesticus* oocytes, BUB1 depletion increased the number of eggs with sister separation (i.e., PSSC), consistent with BUB1’s role in protecting cohesin (*50*). BUB1 depletion in hybrid oocytes also induced PSSC, but importantly, significantly reduced the proportion of eggs with unseparated bivalents. As a complementary approach, we overexpressed BUB1 in *domesticus* oocytes to test if higher BUB1 levels are sufficient to drive the non-separation phenotype (Fig. 3E). Indeed, overexpressed *domesticus* ^EGFP-^BUB1 localized on the kinetochore and to the arm and produced meiosis II eggs with bivalents, supporting our hypothesis. These results are consistent with the idea that the higher BUB1 activity/level in hybrid oocytes drives cohesin over-protection, leading to female sterility.

Since cohesin subunits and their regulators are rapidly evolving, it is possible that cohesion defects occur in other hybrids (*13*, *52*). While searching for other hybrid mouse models with cohesin protection errors, we noticed that hybrid female mice between *Peromyscus maniculatus* and *Peromyscus polionotus* showed significant cohesion defects (i.e., >5 separated sister chromatids) in 5-10% of their meiosis II eggs, while eggs from pure species showed no precocious separation (*53*) (Fig. 4A-C). Visualizing REC8 cohesin in *Peromyscus* meiosis II eggs is technically challenging (*54*), but we found that the proportion of eggs with centromeric REC8 signals was reduced in the hybrid compared to control pure species (Fig. 4D and 4E). As a functional readout of reduced centromeric cohesin, we measured sister-kinetochore distance at metaphase II (Fig. 4E, right graph). Consistent with the REC8 staining result, hybrid eggs showed slightly but significantly increased sister-kinetochore distance compared to pure species. These observations imply a cohesin protection error as in the *domesticus* x *spicilegus* hybrid but in the opposite direction, weakening the protection. To test this idea, we examined the localization of PP2A, which functions with SGO2 to protect cohesin (*49*, *54–57*). We found that PP2A was reduced at the centromere in hybrid oocytes compared to pure species oocytes (Fig. 4F), supporting the idea that hybrid oocytes show weaker cohesin protection. Why do *Peromyscus* hybrid oocytes recruit less PP2A at their pericentromeres? We found that BUB1 kinase and H2A-pT121 levels were significantly lower at the centromere in hybrid oocytes compared to control pure species oocytes (Fig. 4G and 4H). Furthermore, overexpressing *Peromyscus* ^EGFP-^BUB1 partially rescued the cohesin protection defects (Fig. 4B and 4C, no eggs with >5 separated sister chromatids), implying that BUB1 mis-regulation is one of the major causes underlying the defective cohesin protection in *P. maniculatus* x *P. polionotus* hybrid oocytes. *Peromyscus* ^EGFP-^BUB1 overexpression also induced unseparated bivalents in *domesticus* x *spicilegus* oocytes, further confirming its functionality (Fig. 3E). *P. maniculatus* x *P. polionotus* hybrid females do not exhibit substantial fertility defects (*58*, *59*), and therefore, we believe this study has identified an evolving reproductive barrier in this genus.

**Fig. 4.**
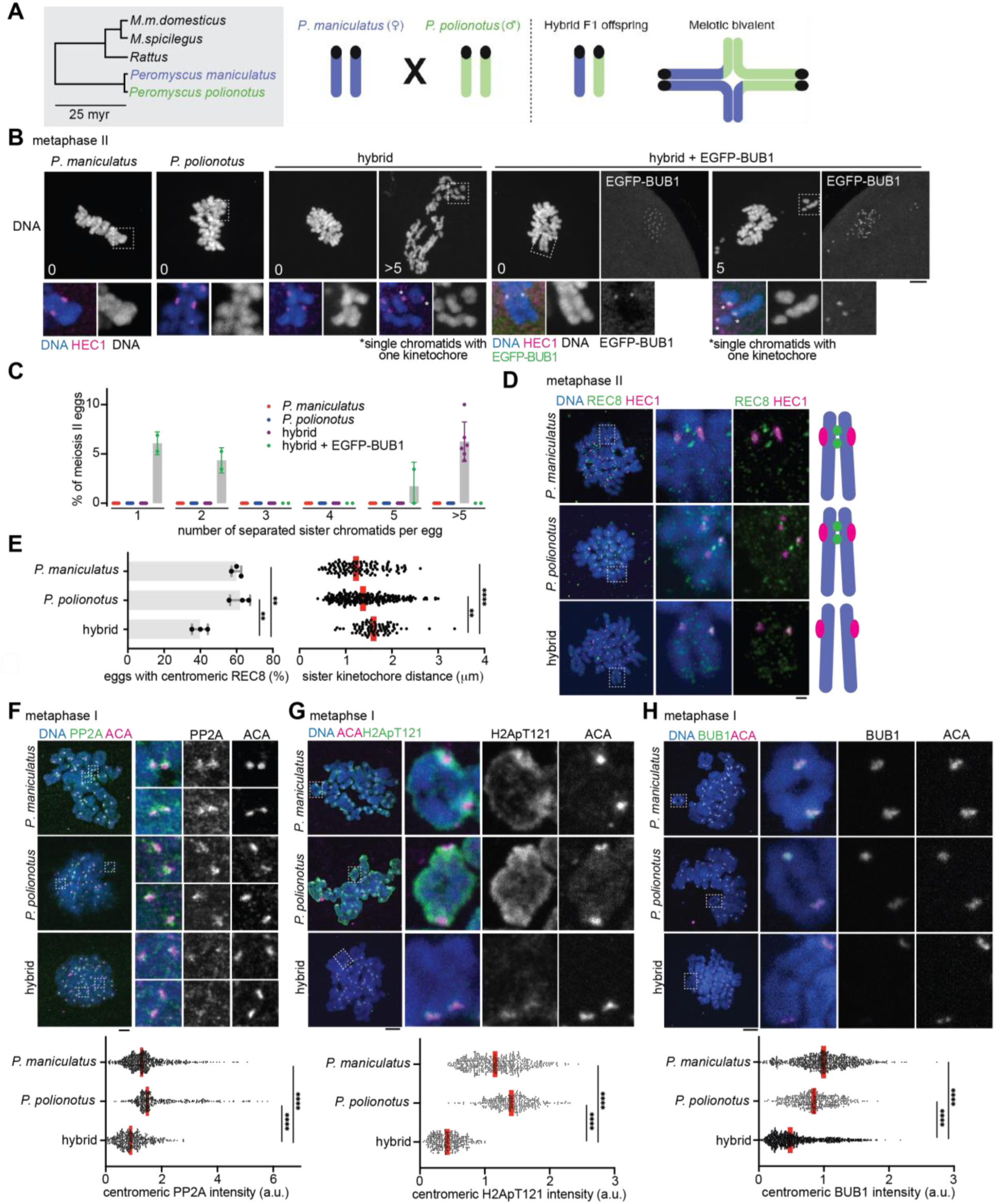
Weaker cohesin protection in *Peromyscus* hybrid oocytes. (**A**) Phylogenetic tree of the *Mus*, *Rattus*, and *Peromyscus* genera (myr, million years) and schematic of the *Peromyscus* hybrid mouse system. (**B**) *P. maniculatus*, *P. polionotus*, and their hybrid oocytes (with or without the EGFP-BUB1 expression) were matured to meiosis II, fixed and stained for HEC1. The numbers in the DNA images indicate the number of split sister chromatids in the egg. (**C**) Graph shows the quantification of the number of separated sister chromatids in each meiosis II egg (n = 162, 171, 147, and 50 eggs for *P. maniculatus*, *P. polionotus*, hybrid, and hybrid + EGFP-BUB1, respectively); each dot represents an independent experiment; bars and error bars represent mean and standard deviation, respectively. (**D**) *P. maniculatus*, *P. polionotus*, and hybrid metaphase II eggs were fixed and stained for REC8 and HEC1. (**E**) Bar graph shows the proportion of meiosis II eggs with centromeric REC8 signals in each genotype (n = 32, 21, and 40 eggs for *P. maniculatus*, *P. polionotus*, and the hybrid); each dot represents an independent experiment; bars and error bars represent mean and standard deviation, respectively; unpaired two-tailed t test was used for statistical analysis. Images from Fig. 4D were used for the quantification. Dot plot is a quantification of sister-kinetochore distance at metaphase II; images from Fig. 4B were used for the quantification; red line, median; Mann-Whitney test was used for statistical analysis. (**F-H**) *P. maniculatus*, *P. polionotus*, and hybrid oocytes were fixed at metaphase I and stained for ACA together with PP2A (f), H2ApT121 (g), or BUB1 (h). The graphs show the quantification of centromeric signal intensities for PP2A (f, n = 678, 403, and 448 centromeres for *P. maniculatus*, *P. polionotus*, and the hybrid), H2ApT121 (g, n = 473, 301, and 322 centromeres for *P. maniculatus*, *P. polionotus*, and the hybrid), and BUB1 (h, n = 805, 394, and 1213 centromeres for *P. maniculatus*, *P. polionotus*, and the hybrid); each dot represents a single centromere; red line, median; Mann-Whitney test was used for statistical analysis. *\*\*P* <0.01, *\*\*\*\*P* <0.0001; scale bars, 5 *µ*m.

This work provides the first cell biological evidence that cohesin protection errors can be a reproductive isolating barrier in mice. While there are examples of hybrid incompatibility that belong to the same category (e.g., oxidative respiration, nuclear trafficking) (*1*), we think this is the first example of recurrent incompatibility in the same molecular pathway (i.e., cohesin protection) identified in hybrid animals from two separate genera.

Defective cohesion due to cohesin loss and weakened protection is a leading cause of reduced fertility especially with advanced maternal age (*32*, *35*, *36*, *60*). Our work suggests that a major cause of human infertility can have a positive role in mammalian speciation. Together with our previous study on condensin dysfunction in *M. m. domesticus* x *M. spretus* hybrid oocytes, SMC complex, the guardian of chromosome structure across the tree of life, may represent a recurrent factor in reproductive isolation.

## Acknowledgments

We thank Alexander E. Kelly for critically reading the manuscript, Stephen S. Taylor, Michael A. Lampson, Kei-Ichiro Ishiguro, and Yoshinori Watanabe for the antibodies, Iain M. Cheeseman for the Separase sensor plasmids, and for the Akera lab members for discussion. Schematics in the figures are created with BioRenders.com.

## Author contributions

Conceptualization: TA

Methodology: WE, BP

Investigation: WE, BP, TA

Funding acquisition: TA

Supervision: TA

Writing – original draft: WE, TA

Writing – review & editing: WE, BP, TA

## Data and materials availability

Dataset required to reproduce the results in this study will be deposited to FigShare upon publication. Materials generated in the current study are available from the corresponding author on reasonable request.

## Supplementary Materials

Materials and Methods

Figs. S1 to S3

References (*60–63*)

## Materials and Methods

### Mouse strains

Mouse strains were purchased from Envigo (NSA, stock# 033 corresponds to CF1, *Mus musculus domesticus*), Jackson Laboratory (C57BL/6J, stock# 000664, *Mus musculus domesticus*, PANCEVO/EiJ, stock# 001384, *Mus spicilegus*), RIKEN BioResource Research Center (ZBN/Ms, stock# RBRC00661, *Mus spicilegus*), and the Peromyscus Genetic Stock Center at the University of South Carolina (*Peromyscus maniculatus bairdii* (BW strain) and *Peromyscus polionotus subgriseus* (PO strain)). CF-1 and C57BL/6J have complementary advantages as *Mus musculus domesticus* strains: CF-1 is an outbred strain with a significantly higher oocyte yield, and C57BL/6J is an inbred strain that efficiently produces hybrid offspring with PANCEVO/EiJ and ZBN/Ms. For most experiments, *Mus musculus domesticus* C57BL/6J females were crossed to *Mus spicilegus* ZBN/Ms males to generate F1 hybrids. The other direction is less efficient in producing offspring, and we specifically used them in fig. S1C. C57BL/6J x PANCEVO/EiJ F1 hybrid females were used for initial characterization of the hybrid (one of the independent experiments in Fig. 1D, 1H, 2A, and 2C). The mouse strain is now discontinued at the Jackson Laboratory, and we have completely switched to ZBN/Ms. *Peromyscus maniculatus* females were crossed to *Peromyscus polionotus* males to produce F1 hybrids, because the other direction does not produce viable hybrid (*61*). All animal experiments were approved by the Animal Care and Use Committee (National Institutes of Health Animal Study Proposal#: H-0327) and were consistent with the National Institutes of Health guidelines.

### Mouse oocyte collection and culture

Fully grown germinal vesicle (GV)-intact prophase I oocytes were harvested from 6- to 10- week-old female mice in the M2 media (Sigma, cat# M7167) supplemented with 5 µM milrinone (Sigma, cat# 475840) to prevent meiotic resumption (*62*). After the oocyte collection, oocytes were transferred to the M16 media (Millipore, cat# M7292) containing 5 µM milrinone covered with paraffin oil (Nacalai, cat# NC1506764) and incubated at 37°C in a humidified atmosphere of 5% CO_2_ in air. To induce meiotic resumption, milrinone was washed out, and oocytes that did not undergo the nuclear envelope breakdown (NEBD) within 1.5 h after the milrinone washout were removed from the culture. For the *in situ* chromosome counting assay in fig. S2, oocytes were matured for 13 h after NEBD and subsequently treated with 100 µM Monastrol (Millipore, cat# 475879) in the organ culture dish (Falcon, cat# 353037) for 2 h 15 min prior to the fixation (*62*).

### Plasmid construction and cRNA synthesis

Coding sequences of *M. m. domesticus* and *Peromyscus maniculatus bairdii* (BW strain) BUB1 were subcloned into the plasmid *In Vitro* Transcription (pIVT) vector with an N-terminal EGFP (*63*). The plasmids were linearized with Aat II and used as DNA templates to synthetize cRNA (see below). The Separase sensor construct was a generous gift from Dr. Iain M. Cheeseman (*47*). The following primers: 5’GAATTAATACGACTCACTATAGGCCGGCGCCACCATGCCAGAG3’ and 5’GCCTCCAAAAAAGCCTCCTCACTACTTCTGGAATAGCTCAGAG3’ were used for PCR amplification to fuse the T7 promoter sequence to the 5’ end of the sensor construct template DNA. cRNA were synthesized using the template DNA and T7 mMessage mMachine kit (Ambion, cat# AM1340) and purified using the MEGAclear kit (ThermoFisher, cat# AM1908).

### Oocyte microinjection and electroporation

GV-intact prophase I oocytes were microinjected with ∼5 pl of cRNA or antibodies in M2 containing 5 µM milrinone, using a micromanipulator TransferMan 4r and FemtoJet 4i (Eppendorf). cRNA used for microinjections were *Egfp-Bub1* (*M. m. domesticus* or *P. maniculatus bairdii* BUB1 fused with EGFP at the N-terminus, 1184 and 950 ng/µl, respectively), *mCherry-Trim21* (Addgene cat# 105522, *M. musculus domesticus* TRIM21 fused with mCherry at the C-terminus, 1500 or 3000 ng/µl for BUB1 and REC8 TrimAway, respectively), and *hH2B-mScarlet-hRad21-mNeonGreen* (Separase sensor, pNM853, human H2B fused with human Rad21(142-476 a.a.) at the C-terminus with the Rad21 fragment flanked by two fluorescent proteins, mScarlet and mNeonGreen, 750 ng/µl). Following microinjections, oocytes were maintained at prophase I in M16 supplemented with 5 µM milrinone overnight to allow protein expression. To TrimAway BUB1, GV oocytes were microinjected with *mCherry- Trim21* cRNA together with normal goat IgG (EMD Millipore, cat# N102) or anti-BUB1 antibody (generous gift from Dr. Stephen S. Taylor). The oocytes were transferred to M16 supplemented with 5 µM milrinone for 2 h to recover and then transferred to M16 for overnight maturation to metaphase II. To TrimAway REC8, GV oocytes were microinjected with *mCherry-Trim21* cRNA. The oocytes were matured to metaphase II overnight in M16, microinjected with anti-REC8 antibody (generous gift from Dr. Michael A. Lampson) using the piezoXpert (Eppendorf), and incubated in M16 for additional 3 h prior fixation. For SGO2 RNAi, GV oocytes were electroporated with control (Stealth RNAi siRNA Negative Control, Med GC, Invitrogen, cat# 12935300) and SGO2 *siRNA* (custom RNAi from Invitrogen, ggataaagacttcccaggaacttta (*48*)) at 200 nM using NEPA21 Super electroporator Type II (NEPA GENE). The oocytes were transferred to M16 supplemented with 5 µM milrinone, cultured for 24 h for efficient knock down, and transferred to M16 for overnight maturation to metaphase II prior to fixation.

### Immunostaining of whole oocytes and chromosome spreads

Meiosis I oocytes and meiosis II eggs were fixed at metaphase I (7 h from NEBD) or at metaphase II (16 h from NEBD) in freshly prepared 2% paraformaldehyde (Electron Microscopy Sciences, cat# 15710) in 1x PBS (Quality Biological, cat# 119-069-101CS) with 0.1% Triton X- 100 (Millipore, cat# TX1568-1) for 20 min at room temperature (RT), permeabilized in 1x PBS with 0.1% Triton X-100 for 15 min at RT, placed in the blocking solution (0.3% BSA (Fisher bioreagents, cat# BP1600-100) and 0.01% Tween-20 (ThermoFisher, cat# J20605-AP) in 1x PBS) overnight at 4°C, incubated for 2 h with primary antibodies at RT, washed three times for 10 min with the blocking solution, incubated for 1 h with secondary antibodies at RT, washed three times for 10 min in the blocking solution, and mounted on microscope slides with the Antifade Mounting Medium with DAPI (Vector Laboratories, cat# H-1200).

For chromosome spreads, zona pellucida was removed from oocytes/eggs, and the oocytes/eggs were fixed with 1% paraformaldehyde, 0.15% Triton X-100, and 3 mM DTT (Sigma, cat# 43815) at metaphase I (7 h from NEBD) or at metaphase II (16 h from NEBD)

The following primary antibodies were used: rabbit anti-mouse REC8 antibody (1:500, gift from Dr. Michael A. Lampson), mouse anti-human HEC1 antibody (1:100, Santa Cruz, cat# sc- 515550), rabbit anti-mouse SGO2 antibody (1:50, gift from Drs. Kei-Ichiro Ishiguro and Yoshinori Watanabe), CREST human autoantibody against centromere (1:100, Immunovision, cat# HCT-0100), sheep polyclonal anti human-BUB1 antibody, SB1.3 (1:100, gift from Dr. Stephen S. Taylor), rabbit anti-human Topoisomerase II (1:100, Abcam, cat# ab109524), rabbit anti-H2ApT120 antibody (1:2500, Active motif, cat# 39391), goat anti-GFP antibody (1:100, Rockland, 600-101-215M), and mouse anti-human PP2A C subunit (1:100, EMD Millipore, cat# 05-421-AF488).

Secondary antibodies were Alexa Fluor 488–conjugated donkey anti-mouse (1:500, Invitrogen, cat# A21202) or donkey anti-goat (1:500, Invitrogen, cat# A11057), Alexa Fluor 568–conjugated goat anti-rabbit (1:500, Invitrogen, cat# A10042), or Alexa Fluor 647– conjugated goat anti-human (1:500, Invitrogen, cat# A21445).

### Granulosa cell immunostaining

The procedure for isolating and culturing ovarian granulosa cells has been described previously (*64*). Briefly, after euthanizing the mice, their ovaries were collected and rinsed three times with M2 media (Sigma-Aldrich, cat# M7167) to remove any adherent fat tissue. The ovaries were then mechanically disrupted to release oocytes and granulosa cells. Following the collection of oocytes, the remaining granulosa cells were collected into a 15 ml tube and allowed to settle at the bottom for 5-10 minutes. The supernatant was discarded to remove blood cells, and the granulosa cells were then centrifuged at 500 x g for 5 minutes. The cells were washed extensively with DMEM high glucose GlutaMAX media (Gibco, cat# 10566-016) supplemented with 1x Antibiotic-Antimycotic (Gibco, cat# 15240062). Cells were dispersed by pipetting, washed for two additional times, seeded on glass bottom chamber slides (Lab-Tek, cat# 155411) with DMEM supplemented with 10% FBS (Gibco, cat# A3160501) and 1x Antibiotic-Antimycotic, and cultured until they reach 50% confluency in a humidified atmosphere containing 5% CO2 at 37°C. After 24 h, the medium was replaced with fresh media of the same type to continue the primary culture. Mitotic cells were enriched by adding 1 μM nocodazole (Sigma-Aldrich, cat# 487929-10MG-M) to the medium and the cells were cultured for 10 h before proceeding to immunostaining (see above).

### Confocal microscopy

Fixed oocytes, eggs, chromosome spreads and granulosa cells were imaged with a microscope (Eclipse Ti; Nikon) equipped with 100x / 1.40 NA oil-immersion objective lens, CSU-W1 spinning disk confocal scanner (Yokogawa), ORCA Fusion Digital CMOS camera (Hamamatsu Photonics), and 405, 488, 561 and 640 nm laser lines controlled by the NIS-Elements imaging software (Nikon). Confocal images were acquired as Z-stacks at 0.3 µm intervals. For live imaging, oocytes were placed into 3 µl drops of M16 covered with paraffin oil in a glass-bottom tissue culture dish (fluoroDish, cat# FD35-100) in a stage top incubator (Tokai Hit) to maintain 37°C and 5% CO_2_. Time-lapse images were collected with a microscope (Eclipse Ti2-E; Nikon) equipped with the 20x / 0.75 NA objective (Fig. 1D and 2A and fig. S1A and S1B), CSU-W1 spinning disk confocal scanner (Yokogawa), ORCA Fusion Digital CMOS camera (Hamamatsu Photonics), and 405, 488, 561 and 640 nm laser lines controlled by the NIS-Elements imaging software (Nikon). Confocal images were collected as Z-stacks at 3 µm intervals to visualize all the chromosomes. Images are displayed as maximum intensity Z-projections in the figures.

### Image analysis

Fiji/ImageJ (NIH) was used to analyze all the images. In general, optical slices containing chromosomes were added to produce a sum intensity Z-projection for pixel intensity quantifications. For line scans of SGO2, BUB1, and H2ApT121 signal intensities (Fig. 2B, 3A and C), lines (width: 11 pixels) were drawn from the centromere towards the chromosome arm, and signal intensities were averaged over multiple chromosomes after subtracting background signals, obtained near the chromosome. To quantify chromosomal H2ApT121 signal intensities in oocytes (Fig. 3B), masking images were created using DAPI staining images to specifically measure the signal intensities on the chromosome. The signal intensity was integrated over each slice after subtracting the background, obtained near the chromosomes. For the quantification of the Separase sensor cleavage (Fig. 2A), masking images were created using mScarlet images to specifically measure signal intensities of mNeonGreen and mScarlet on the chromosomes from metaphase I to anaphase I. Signal intensities were integrated over each slice after subtracting the background, obtained near the chromosomes. The mNeonGreen / mScarlet ratio was calculated for each timepoint. For the *in situ* chromosome counting assay (Fig. S2), the number of chromosomes were counted in metaphase II eggs using the DAPI and TOP2A (centromere) signals. To specifically quantify centromeric signal intensities (PP2A, BUB1, and H2A-pT121 in Fig. 4F-H), ellipses were delineated around the centromere (based on the ACA staining) on each chromosome. Signal intensities were then quantified within each ellipse after the background signal subtraction. Representative images in the figures are maximum intensity Z-projections unless specified in the figure legend.

### Statistics and reproducibility

Data points were pooled from at least two independent experiments. Statistical analyses were performed using Microsoft Excel and GraphPad Prism 10. Scattered plots and line and bar graphs were created with GraphPad Prism 10. Mann-Whitney test and unpaired two-tailed t test were used for statistical analysis unless specified in the figure legend, and a value of *P* < 0.05 was considered significant.

**Fig. S1. Cohesin is maintained along the chromosome arms in hybrid oocytes.** (**A**) Mis-segregation in *domesticus* x *spicilegus* oocytes is associated with cytokinetic failures. Images from Fig. 1D were analyzed to quantify the proportion of oocytes with cytokinetic failure. (**B**) Examples of non-separated bivalent chromosomes captured by live-imaging. The images are from the same time-lapse imaging dataset from Fig. 2A. (**C**) Chromosome spreads were performed at metaphase II using hybrid oocytes derived from both cross directions (i.e., *domesticus* x *spicilegus* and *spicilegus* x *domesticus*) and stained for HEC1 and REC8. Graph shows the quantification of the number of bivalents per egg (n = 13 and 13 meiosis II eggs for *domesticus* x *spicilegus* and *spicilegus* x *domesticus*, respectively); note that the data for *domesticus* x *spicilegus* is from Fig. 1G; each dot represents a single egg; red line, mean; unpaired two-tailed t test was used for statistical analysis; \*\**\*\*P* <0.0001.

**Fig. S2. Hybrid meiosis II eggs from older female mice carry less bivalents.** Hybrid oocytes from young (3-month old) and aged (12-month old) mice were matured and fixed at metaphase II and stained with TOP2A (centromere). Graph quantifies the number of bivalents in each meiosis II egg (n = 21 and 36 eggs from young ang aged mice, respectively); each dot in the graph represents a single egg; red line, median; Mann-Whitney test was used for statistical analysis; *\*\*P* <0.01.

**Fig. S3. Sister kinetochores are normally split in unseparated bivalents.** Chromosome spreads were performed at metaphase I (*domesticus*) and metaphase II (*domesticus* and the hybrid) and stained for HEC1. Graph shows the quantification of the percentage of chromosomes with separated sister-kinetochores (i.e., inter-kinetochore distance larger than 1.2 µm) (n = 104, 100, and 89 chromosomes for *domesticus* metaphase I, *domesticus* metaphase II, and hybrid metaphase II).

