## Supplementary figures and images for "Hybrid female sterility due to cohesin protection errors in oocytes"

### figures S1-S3

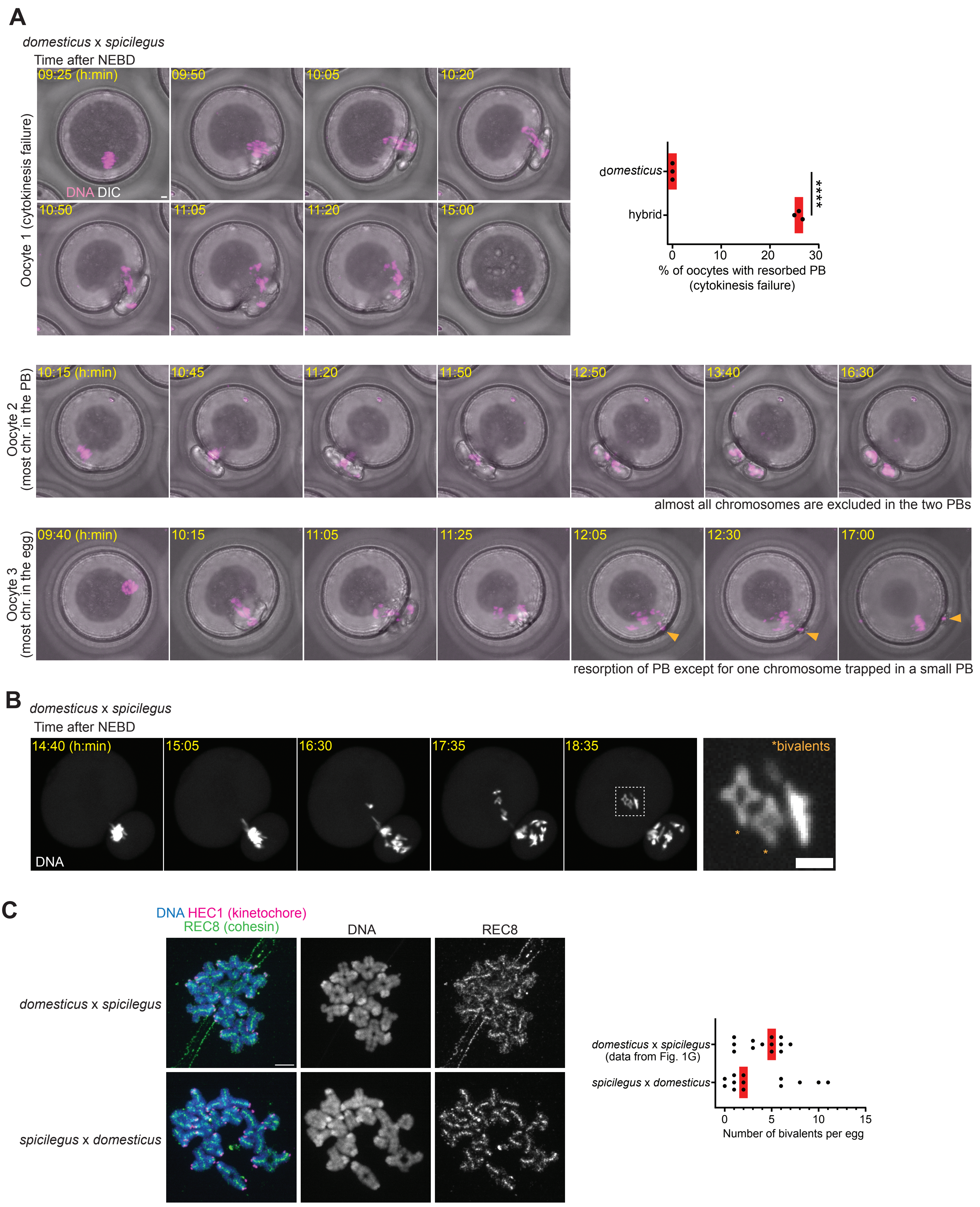

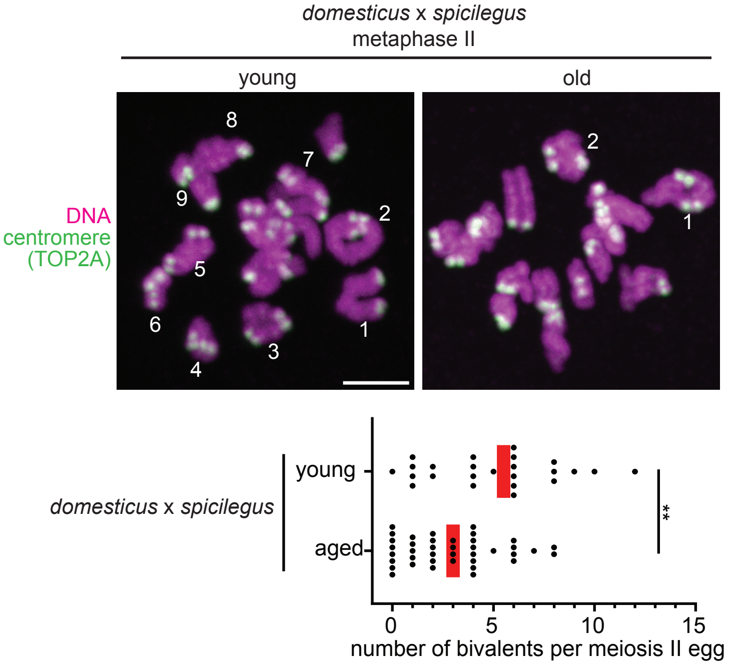

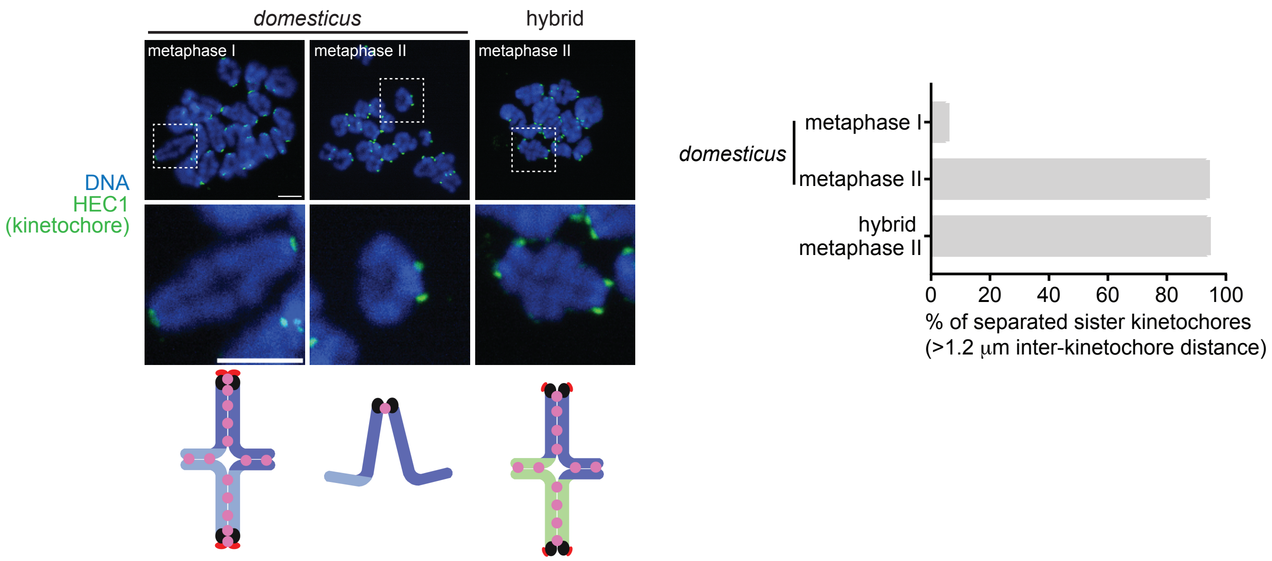
